# A comprehensive characterization of rhythmic spiking activity in the rat ventral striatum

**DOI:** 10.1101/617233

**Authors:** Matthijs A. A. van der Meer, Jimmie M. Gmaz, J. Eric Carmichael

## Abstract

The ventral striatum (vStr) is anatomically interconnected with brain structures that exhibit prominent rhythmic activity, suggesting that oscillations in ventral striatal activity are potentially informative about systems-level interactions between these structures. However, rhythmic activity in ventral striatal neurons during behavior has only been characterized piecemeal, with individual studies focusing on a single cell type or frequency band. We performed a comprehensive analysis of (1) rhythmic activity in vStr neurons without reference to the local field potential, and (2) average as well as time-resolved spike-field relationships. Spike train rhythmicity tended to be limited to low frequencies such as delta and theta, whereas spike-field relationships were seen across a broad spectrum of frequencies, with about 90% of neurons showing spike-field locking to at least one rhythm. Using a novel time-resolved generalized linear model approach, we further show that the contribution of local field potential (LFP) phase to spike timing is dynamic over time, and enhanced by the inclusion of the LFP from the hippocampus – a new measure of inter-area coupling. These results provide a foundation for a more accurate interpretation of the ventral striatal LFP, suggest the possibility of an oscillatory taxonomy of ventral striatal neurons, and provide a starting point for understanding how rhythmic activity links cell-, circuit-, and systems-level phenomena in the ventral striatum.

**Significance Statement:** Oscillations in neural activity are ubiquitous in the brain, readily accessible in the clinic and the lab, and shared by humans and animals to facilitate translational work. The ventral striatum (vStr) is a promising target structure for such a rhythmic activity perspective, not in the least because its local field potential (LFP) shows prominent task-related oscillations across a range of frequencies. However, recent work has shown that major components of the vStr LFP are in fact generated elsewhere in the brain, raising the question of how the LFP relates to local spiking activity. Unlike previous studies that focused on a specific cell type or frequency band of interest, we characterize rhythmic activity across a full range range of frequencies and cell types, and include novel analyses appropriate for a non-local LFP. Our results provide a foundation for more accurate interpretation of the vStr LFP and a starting point for an oscillatory taxonomy of vStr neurons.

## Introduction

Rhythmic fluctuations in neural activity are pervasive throughout the brain, and the study of such oscillations have provided a powerful window on the dynamic nature of neural computation (Engel et al., 2001; Buzsáki, 2006; Womelsdorf et al., 2014). For instance, in the hippocampal formation, prominent theories of navigation propose that velocity-controlled oscillators are a core component of a path integration system (Burgess and O’Keefe, 2011). In the cortex, top-down and bottom-up communication is associated with beta and theta-gamma frequencies, respectively (Bastos et al., 2015). More generally, rhythmic fluctuations in excitability create opportunities for selective gain control (Fries, 2015). Even for those who are skeptical whether oscillations have mechanistic relevance, numerous studies have shown oscillations can provide a useful readout of neural activity. For instance, pre-stimulus rhythmic activity can be used to predict responses to near-threshold stimuli (Lakatos et al., 2008; Busch et al., 2009; Spaak et al., 2014). Local field potentials oscillations can be used to improve the effectiveness of deep brain stimulation (Rosin et al., 2011; Priori et al., 2013). Numerous pathological brain states are associated with altered brain rhythms, including at the prodromal stage (Jenkinson and Brown, 2011; Uhlhaas and Singer, 2015; Tada et al., 2016). Thus, possible mechanistic relevance aside, there are many practical reasons to be interested in rhythmic neural activity.

In the ventral striatum (vStr), a brain structure involved in the motivational control of behavior, several conditions are in place to suggest that oscillations can usefully inform studies of its function. First, the vStr LFP shows a full spectrum of rhythmic activity, including prominent gamma-band oscillations but also delta, theta, and beta-band activity, which are modulated in association with behavior and task variables and relate to local spiking activity (Leung and Yim, 1993; van der Meer and Redish, 2009; Berke, 2009; Kalenscher et al., 2010; Howe et al., 2011; Donnelly et al., 2014; Dejean et al., 2017; Dwiel et al., 2019). Manipulations of the dopaminergic and endocannabinoid system modify vStr LFP oscillations (Berke, 2009; Lemaire et al., 2012; Morra et al., 2012) and drug and disease states are associated with altered striatal LFPs (Dejean et al., 2017; Naze et al., 2018; Wu et al., 2018; Hultman et al., 2018). Striatal neurons show intrinsic rhythmic activity and frequency-specific resonance (Bracci et al., 2003; Taverna et al., 2007; Beatty et al., 2015). Finally, the vStr LFP dynamically synchronizes with LFPs in anatomically related structures such as the (pre)frontal cortex and the hippocampus (Gruber et al., 2009; van der Meer and Redish, 2011; Catanese et al., 2016; Lansink et al., 2016). Taken together with the known convergence of these multiple rhythmic inputs onto vStr neurons, these observations suggest considerable fundamental and translational research potential of oscillations in the vStr.

However, an obstacle to realizing this apparent promise is that major components of the ventral striatal LFP turn out to be non-local: LFP gamma oscillations are volume-conducted from the piriform cortex (Carmichael et al., 2017), while LFP theta oscillations are volume-conducted from the septum (Lalla et al., 2017). Nevertheless, we and others have found consistent spike-field relationships in the vStr, which likely result from direct inputs and efference copy associated with these areas (Karalis and Sirota, 2018). This indirect relationship between the vStr LFP and local spiking raises the possibility that the vStr LFP can range from completely dissociated from spiking (no spike-field relationship; e.g. if nearby structures generating the LFP are not influencing vStr spiking) to strongly related to spiking (e.g. if a LFP-generating structure drives striatal spiking). Given this indirect relationship, it is particularly important to understand (1) the rhythmic properties of striatal spiking without reference to the LFP, and (2) under what circumstances the LFP is informative about striatal spiking. In addition, although several studies have characterized spike-field relationships in the striatum, they have done so for specific cell types (e.g. only FSIs: van der Meer and Redish 2009; MSNs: Kalenscher et al. 2010; or specific frequencies, theta: van der Meer and Redish 2011, beta and gamma: Howe et al. 2011, gamma: van der Meer and Redish 2009, Kalenscher et al. 2010). Thus we lack a cohesive view of spike-field relationships across cell types and frequencies, and of how such relationships relate to local rhythmic spiking without reference to the field potential.

To address these issues, we performed a number of analyses, applied to neurons recorded extracellularly from the ventral striatum as rats performed a modified T-maze task. First, we computed *spike spectra* to characterize spike train rhythmicity without reference to the LFP. Next, we characterized *spike-field relationships* across all frequencies for both putative MSNs and FSIs, including their preferred phase distributions. Finally, we introduce a novel time-resolved application of *generalized linear models (GLMs)* to determine the improvement over baseline models in dynamically predicting spike times afforded by different LFP variables, including those from the hippocampus, an input to the ventral striatum.

## Materials and Methods

### Data

This study uses the combined data previously published as (van der Meer and Redish, 2009, 2011). Briefly, male Brown Norway/Fisher-344 hybrid rats (n = 11) performed variations of a continuous T-maze task. Daily recording sessions consisted of a pre-task rest epoch, a task epoch, and a post-task rest epoch. Only task epoch data was used, consisting of a number of *trials* (defined as laps on the T-maze). Correct trials yielded reward at two reward sites on the chosen arm of the T-maze. Ventral striatal local field potentials and multiple single units were acquired from all subjects; some subjects (n = 7) additionally had recording electrodes in the dorsal CA1 area of the hippocampus, of which only the fissure LFP was used for analysis. Single units were labeled as putative medium spiny neurons (MSNs), putative fast-spiking interneurons (FSIs) and other, following the method in van der Meer and Redish 2009, based on waveform properties and average firing rate (Berke et al., 2004).

### Data analysis overview

This study contains three main analyses: (1) spike spectra, designed to characterize spike train rhythmicity without reference to the LFP, (2) spike-field relationships, as measured by the pairwise phase consistency (PPC, Vinck et al. 2010) and spike-triggered averages, and (3) generalized linear models that aim to predict spike times, designed to reveal how much LFP variables improve spike timing prediction above and beyond prediction based on non-LFP factors alone. Analyses (1) and (2) are illustrated schematically in Figure 1, and analysis (3) is illustrated in Figure 2.

**Figure 1:**
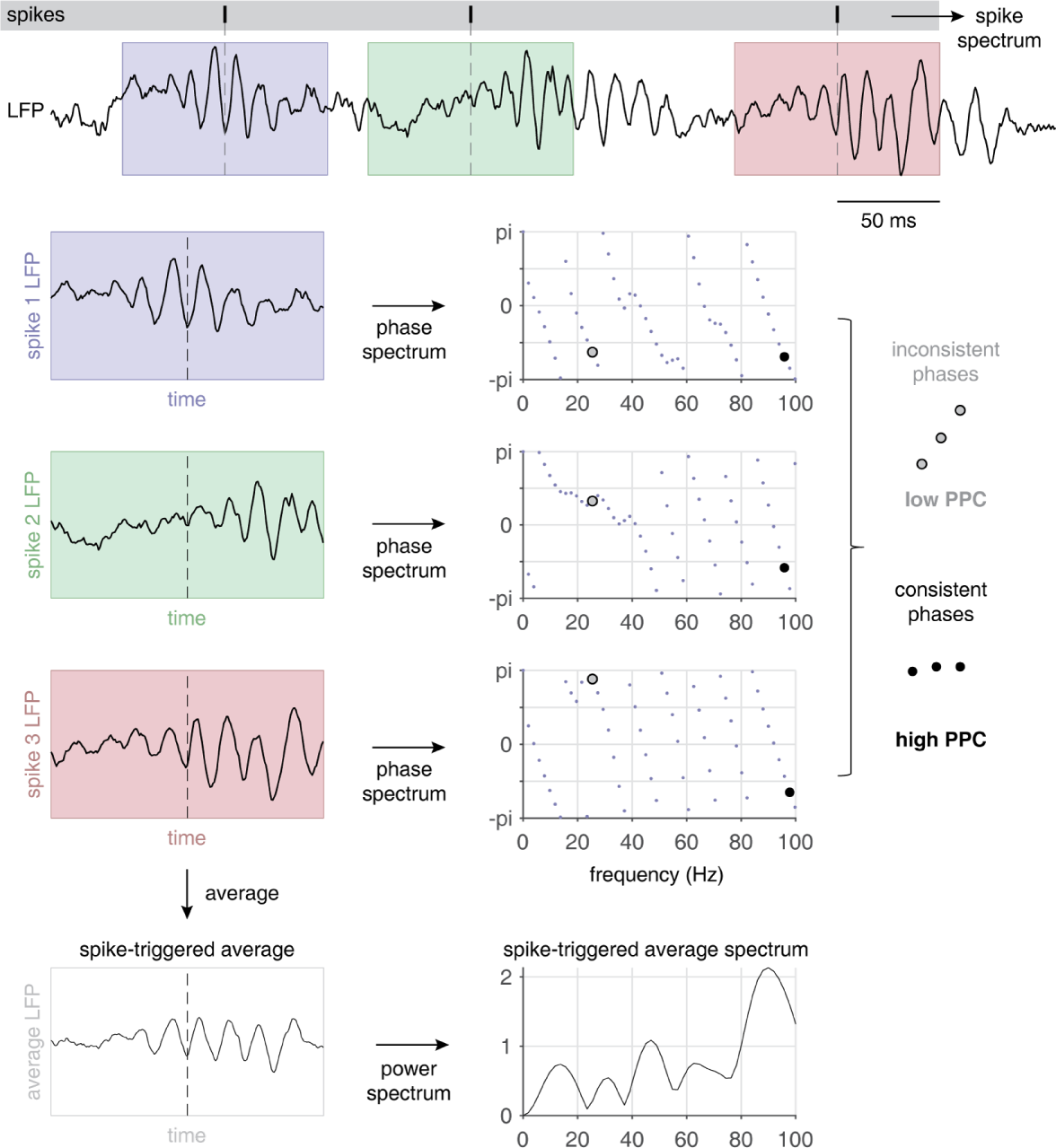
Schematic illustration of the spike spectrum, pairwise phase consistency (PPC) and spike-triggered average (STA) analyses. Top panel shows an example 500 ms trace of a ventral striatal LFP, along with a spike train (vertical tickmarks on gray background) shown on the same time base. The spike spectrum for a given cell is computed based on spike times alone (no reference to the LFP). PPC and STA both start by taking spike-triggered snippets of the LFP, shown as blue, green and red windows for the three spikes in this example. To obtain the PPC spectrum, the phase spectrum (angle of the Fourier transform) is computed for each spike-triggered LFP. For each frequency, PPC quantifies how consistent the phases are across all (pairs of) spike-triggered LFPs. In this example, spike phases are inconsistent at 23 Hz (open circles) but consistent at 97 Hz (filled circles). To obtain the STA spectrum, the spike-triggered LFPs are averaged and then the power spectrum computed.

**Figure 2:**
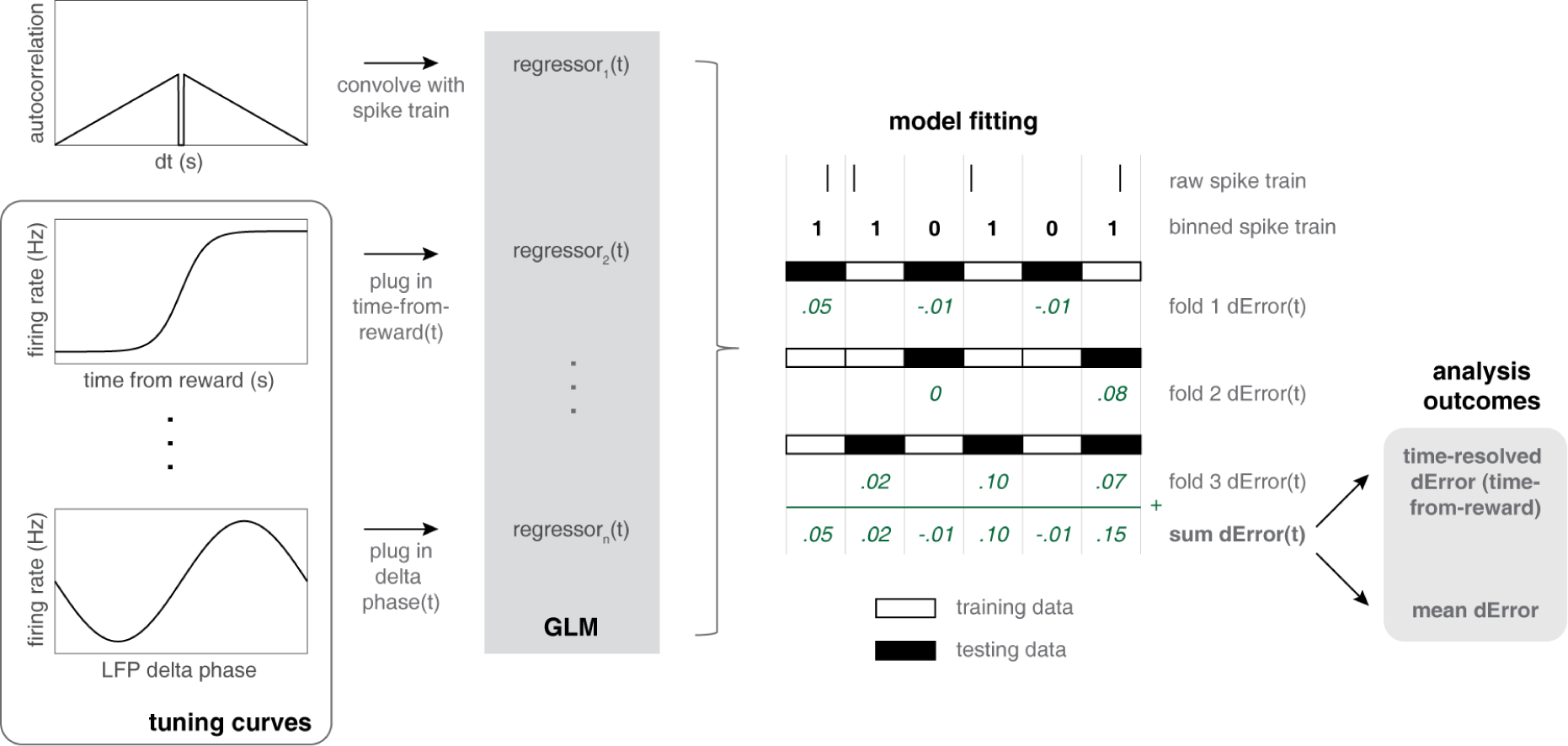
Schematic illustration of the time-resolved generalized linear model (GLM) approach. For each spike train, binarized into 1 ms bins, a set of regressors is constructed, starting with a conditional intensity function (cif) based on the cell’s autocorrelation (top left). Intuitively, this regressor captures the known dependencies in the cell’s spike train to estimate predicted firing rate at time *t* + *dt* given that there was a spike at time *t*. Next, all other regressors in the GLM were constructed using a two-step process. For each variable of interest, a *tuning curve* is estimated, which shows average firing rate as a function of the relevant variable (e.g. time from reward, or the delta-band phase of the LFP). For each time bin in the GLM, this tuning curve can be used to look up the predicted firing rate based on the value of the relevant variable at that time (e.g. delta phase at time *t*). Next, for each cross-validation fold, the binned spike train is divided into training (clear bins) and testing sets (black bins), and the squared error for each testing bin computed for all candidate models. Specifically, the difference between a baseline model’s error and the error of alternative models (*dError*, typically including one or more features of the local field potential) is stored for each bin and each cross-validation fold. Then, this error is averaged *across folds* to yield a difference in model fits as a function of time, which can either be averaged *over time* to yield an overall measure of model performance relative to baseline, or plotted relative to time of reward delivery.

### Data inclusion criteria

The data in each session were first restricted to a ± 5 s window centered on the time of reward delivery at the first reward site. This window was chosen because it contains the major behavioral states associated with vStr LFP rhythms: theta during reward approach, beta during trial completion, and delta/gamma during reward consumption and rest. In order to provide reliable estimates of rhythmic activity and phase locking, any unit firing less than 100 spikes in the remaining time was excluded from analysis. Because estimates of spike train rhythmicity and spike-field locking are sensitive to cluster quality, we excluded cells with isolation distance smaller than 20, and cells with a L-ratio larger than 0.1 (Schmitzer-Torbert et al., 2005). Finally, to reduce the possibility that the same cell recorded across days was counted multiple times in the analysis, we excluded possible duplicates, defined as having a waveform correlation (across all tetrode wires) of at least .99 or higher, and a normalized peak difference of .2 or less. Because spike waveforms have significant power even at low frequencies, which can result in artefactual spike-field relationships (Zanos et al., 2010) we excluded all cells recorded from the same tetrode as the LFP used.

### Spike spectra

This analysis computes a multitaper estimate of the frequency content in spike trains. Intuitively, the spike spectrum captures how rhythmic spike times occur at each frequency. It is related to the power spectrum of the spike train’s autocorrelation through the Wiener-Khinchin theorem (Gabbiani and Koch, 1998). To compute spike spectra, we used the `mtspectrumpt` function from the Chronux toolbox (Mitra and Bokil, 2008), with the following parameters: tapers [7 11], *F*_*s*_ = 200.

### Spike-field relationships

Contemporary measures of spike-field relationships are based on the *spike-triggered spectrum (STS)*, i.e. the discrete Fourier transform of a short window of LFP data centered on each spike (Figure 1). Spike-triggered spectra offer the dual advantages of spanning all frequencies (at a resolution defined by the parameters of the spectral estimator used), and being time-resolved (the time of each spike-triggered spectrum is retained). The consistency of the phases across spike-triggered spectra is used to quantify phase locking at each frequency; we use the pairwise phase consistency measure (PPC; Vinck et al. 2010) which is 0 for random phases and 1 for perfect phase locking^1^. To obtain the spike-triggered average (STA) spectrum, spike-triggered LFPs are first averaged and then the power spectrum computed. Note that this is not the same as the PPC, because in PPC all phases are weighted equally regardless of LFP power, whereas in the STA each spike contributes to the average according to LFP power. To compute PPC, we used the FieldTrip toolbox (Oostenveld et al., 2011)’s `ft_spiketriggeredspectrum` and `ft_spiketriggeredspectrum_stat` functions, with a frequency-dependent time window to estimate spectral content: *cfg.toi* = 5*/f*.

We chose PPC as our primary measure of spike-field locking because unlike measures based on mean vector length of spike phases such as phase-locked value (PLV) or Rayleigh’s *r*, PPC does not overestimate phase locking strength for small numbers of spikes, facilitating comparison across multiple data sets and cell types that may have different numbers of spikes (Vinck et al., 2010; Aydore et al., 2013). We elected not to use the corrected PPC measure that rules out history effects by only using pairs of spikes from different trials (Vinck et al., 2011); although our data was structured in trials (the 10-second windows around each reward delivery) we want to facilitate comparison with non-trialified data in the future. In addition, our GLM analysis (described next) provides a more general approach to ruling out possible confounds in estimating spike-field relationships. Nevertheless, for comparison with what some may see as a more intuitive measure of spike-field relationships, we also computed the spike-triggered average (STA) spectrum using the Field-Trip functions `ft_spiketriggeredaverage` and `ft_spectrum` with the same frequency-dependent time window as above.

### Shuffles and statistical significance

To assess the statistical significance of the spike spectra and spike-field locking procedures, we compared the observed values to resampled distributions, obtained from 1000 shuffles of the spike times of each cell. (To speed up this procedure, we pre-computed for each recording session a large pool of spike-triggered spectra based on uniformly distributed spike times, and for each shuffle selected a random subset from this pool of the same size as the number of spikes in the analyzed cell.) The distributions of resampled values thus obtained were used to derive z-scores for the observed data (i.e. how many standard deviations away from the shuffled mean the actual data is). We applied an arbitrary threshold of *p* < 0.05 uncorrected for determining proportions of cells with significant rhythmic activity at each frequency or frequency band.

### GLM analysis

In our final set of analyses, we sought to construct models that, for each cell, attempt to predict the probability of a spike in each 1 ms time bin, given a set of predictor variables (regressors) and a linear model with a logit link function (Truccolo et al., 2005; Sarma et al., 2012; Zhou et al., 2015). This approach affords a few advantages over isolated measures of spike-field relationships: first, by putting all predictor variables on an equal footing, it can address how much *additional* variance is explained by incorporating variables of interest (for spike-field relationship questions, these would be LFP phases at various frequencies) after the effects of other variables are accounted for. If these variables include, for instance, the cell’s autocorrelation function, then history effects can be accounted for. If a cell’s tuning to task variables such as reward receipt is included, then correlations between task variables and LFP phases are accounted for, and so on. Second, the GLM framework provides a way to compare the contributions of LFP phase to other variables that appear to be related to vStr spiking; based on a PPC of 0.01, for instance, it is unclear if this provides a little, or a lot, of predictive power compared to knowing, say, the cell’s tuning to a task variable such as the time relative to reward delivery. Finally, GLMs are an elegant way to progress toward a longer-term goal of being able to account for all vStr activity by incorporating more variables. For instance, if the inclusion of a LFP in a *different* brain structure improves the model, this may indicate effective connectivity between these structures(Wong et al., 2016).

Our specific implementation of the spike train GLM generally follows established methods (Kass et al. 2014; Kramer and Eden 2016; we used the MATLAB function `fitglm`, spike trains were binned at 1 ms). However, we apply two innovations. First, although we used a standard cross-validation approach for model comparison, we kept track of the error at each time bin (instead of compressing the model fit for each fold into a single mean-squared-error number) so that we could later plot model improvement as a function of task variables. This approach, illustrated schematically in Figure 2, makes it possible to plot the difference in error between a baseline model and the target model (typically baseline plus one or more LFP phase variables) as a function of task variables such as the time relative to reward delivery. In this way, this approach can reveal whether the additional predictive value of LFP variables is constant across the task, or is modulated in relation to various task events and behaviors such as reward approach and consumption.

The second GLM innovation lies in the specific way we included various task features such as time to reward delivery, running speed, and position on the track as predictors. Instead of plugging these variables directly into the GLM as regressors, we first constructed *tuning curves* for each neuron in each of these task variables (i.e. average firing rate as a function of task variable value) and used the predicted firing rate from this tuning curve in the GLM (Figure 2, lower left). This is important because typical neurons have nonmonotonic tuning to these variables, such that a linear model would fail to capture their contribution accurately. Consider, for instance, a typical “place cell” which first increases and then decreases its firing as a function of (linearized) location. Raw linearized position would be an ineffective predictor for such a cell, but the tuning curve regressor method computes the expected firing rate for each location and feeds that into the GLM.

GLM regressors used in this way include (1) time to reward (ranging from −5 to 5 seconds, relative to reward delivery), (2) linearized position on the maze, and (3) running speed. Next, we include a conditional intensity function regressor (Truccolo et al., 2005; Rule et al., 2015) based on each cell’s autocorrelation function by convolving the binned spike train with the autocorrelation function (a 1-second window centered on the spike times). Finally, LFP phase features were computed by filtering the data in specific frequency bands of interest (delta: 3–5 Hz, theta: 7–9 Hz, beta: 14–25 Hz, low-gamma: 40–65 Hz, high-gamma: 70–95 Hz) and taking the `angle` of the Hilbert-transformed data.

## Results

We sought to characterize rhythmic activity in single neurons in the rat ventral striatum (vStr) during the performance of a continuous T-maze task (van der Meer and Redish, 2009). Data from 81 daily recording sessions from 4 rats was restricted to a 10-second time window centered on the time at which rats first reached a reward site following correct choice. As we and others have previously shown, reward approach and receipt elicits a dynamic pattern of oscillations in the ventral striatal LFP, including theta oscillations which dominate during approach (van der Meer and Redish, 2011; Lansink et al., 2016; Sjulson et al., 2018), beta oscillations upon cue utilization and trial completion (Howe et al., 2011; Leventhal et al., 2012), and delta and gamma oscillations during reward consumption and immobility more generally (Donnelly et al. 2014; Malhotra et al. 2015; Figure 3a shows a single-session spectrogram illustrating these LFP components). Thus, this time window is known to contain the major LFP rhythms whose associated rhythmic spiking patterns we aim to characterize.

**Figure 3:**
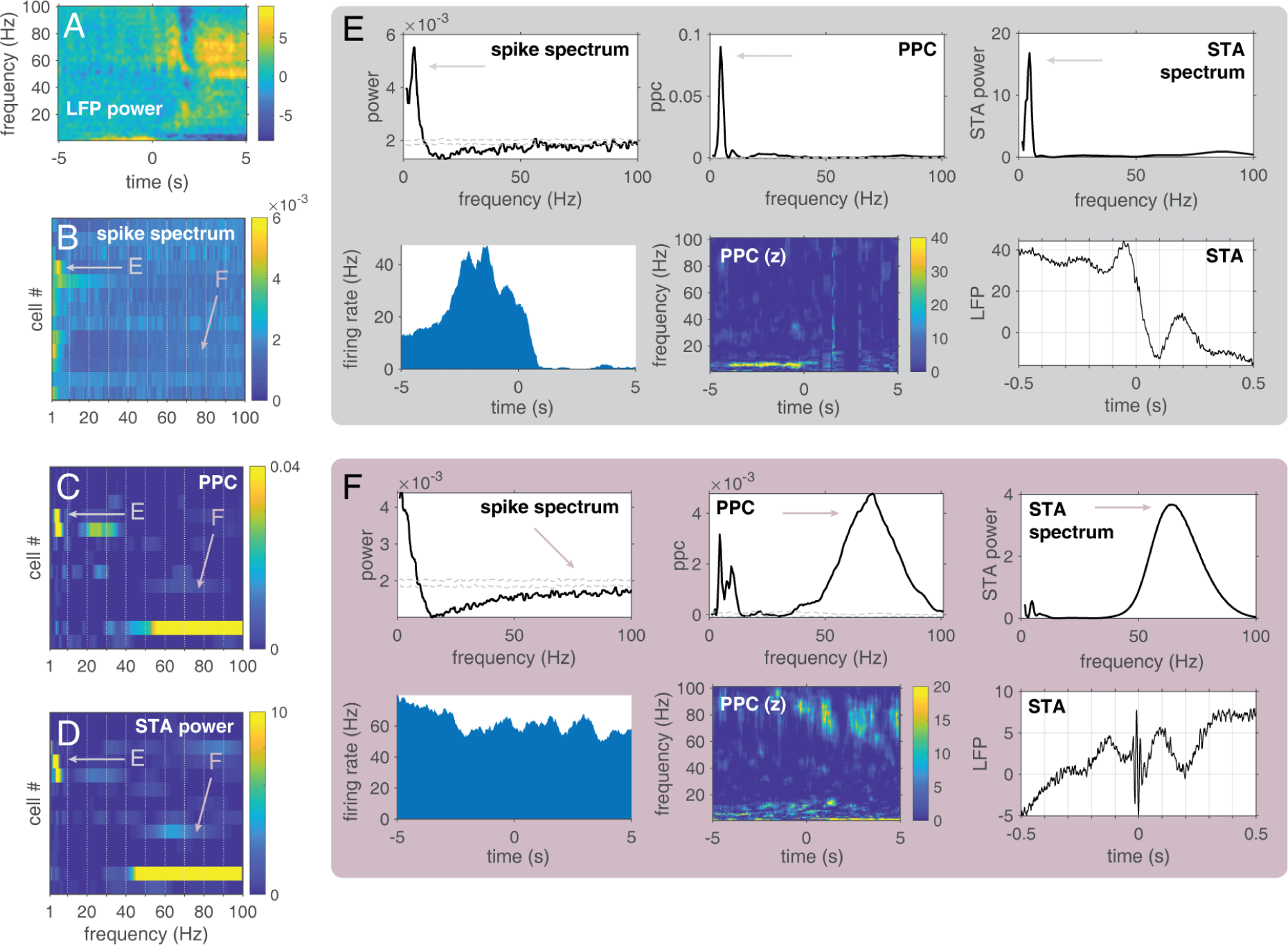
Cell-specific and dynamic rhythmic activity in ventral striatal neurons recorded during a representative single session. **A**: LFP spectrogram, showing low frequencies dominating during reward approach (before time 0, which indicates reward delivery) and higher frequencies prominent following reward delivery. **B**: Spike spectra, indicating rhythmic power at each frequency for 13 simultaneously recorded cells. **E** and **F** indicate cells highlighted in detail in the corresponding panels. Pairwise phase consistency (PPC, **C**) and power of the spike-triggered average (STA, **D**), both measure of spike-field locking, are shown for the same cells. **E, F**: Detailed characterization of two example cells. Cell **E** shows consistent peaks at approximately 4.5 Hz in the spike spectrum (top left), PPC (top center) and STA (top and bottom right). Time-resolved PPC (bottom center) indicates this phase locking occurred specifically prior to reward delivery, associated with a “ramping” activity pattern (bottom left shows the peri-event time histogram of firing rate). Cell **F** does not exhibit clear peaks in the spike spectrum, but shows phase locking at several frequencies. Dashed lines in the spike spectrum and PPC plots indicate ± 1 standard deviation around a shuffled mean, obtained by randomly permuting spike times (see *Materials and Methods* for details).

Our initial analysis of rhythmic spiking focuses on (1) the spike spectrum, which describes the frequency content of spiking without reference to the LFP, and (2) two related measures of spike-field relationships: the pairwise phase consistency (PPC) and the power spectrum of the spike-triggered average (STA). The procedures used to obtain these measures are illustrated schematically in Figure 1, and the values obtained for all neurons in a single example session are shown in Figure 3b-d. Inspection of these panels suggests that for PPC and STA in particular, specific frequency ranges are associated with clearly elevated values in individual cells. For instance, the cell highlighted as “E” in each of the panels in Figure 3b-d shows a bright yellow patch at about 4.5 Hz, indicating strong phase locking to the LFP at that frequency, as well as strong rhythmic spiking^2^. The cell highlighted as “F” in these same panels, in contrast, does not show any elevated values in that same frequency range, but instead has elevated PPC and STA power in the gamma band (peaking around 65–70 Hz in this case). This cell does not appear to show corresponding spike spectrum power at that same frequency (note lack of elevation at the “F” arrow in Figure 3b). Further cells not highlighted explicitly also show clear patches of increased PPC and STA power, such as the cell directly below the “E” arrow, and the prominent gamma-band cell at the second row from the bottom.

To facilitate the interpretation of panels 3b-d, we next illustrate in detail the different measures for the two example cells in panels E and F. Cell E (Figure 3e), a putative medium spiny neuron (MSN), shows a consistent 4.5 Hz peak in the spike spectrum (top left), PPC (top middle), and STA (top right). This cell showed a ramp-like firing rate pattern on the task, as indicated by the peri-event time histogram on the bottom left. A time-resolved, z-scored PPC spectrum (bottom middle) shows the clear 4.5 Hz phase locking as a horizontal streak leading up to time 0, the time of reward delivery. The cell in Figure 3f, a putative fast-spiking interneuron, showed no obvious changes in firing rate during the 10-s window (bottom left). As indicated by the PPC and STA plots, this neuron locked to several frequencies in the LFP, most prominently the gamma band (arrows) and to a lesser extent, 4 and 8 Hz noticeable in the PPC spectrum in particular. As noted previously, this phase locking was not associated with clear peaks in the spike spectrum (top left); an intuition for how this can happen is that such cells do not tend to spike on each cycle of the LFP oscillation (explaining the lack of spike spectrum peaks), but when they do spike, there is a LFP phase preference (explaining phase locking). The time-resolved PPC spectrum (bottom middle) indicates that gamma phase locking waxes and wanes dynamically, without any obvious relationships to either firing rate (compare the PETH in the bottom left panel) or to overall LFP power (Figure 3a). Thus, different simultaneously recorded cells show different rhythmic activity profiles.

### Analysis of rhythmic activity across the population

Next, we applied these analyses to the full population of recorded cells. Figure 4 shows the results for putative medium spiny neurons (MSNs, 579 cells analyzed that passed inclusion criteria). The unnormalized **spike spectra** were either relatively flat, or showed increased power at low frequencies, without clear increases at specific frequencies (Figure 4a, left). Accordingly, the average spike spectrum (left panel inset) shows no obvious peaks, other than a hint of a local increase at around 9 Hz (theta). When normalizing the spike spectra by z-scoring relative to spectra obtained from a shuffled distribution, a similar pattern appears, consisting of a strong emphasis on low frequency power with a small theta peak superimposed (Figure 4a, right panel). Thus, although examples of cells with rhythmic spiking at specific frequencies can be found (e.g. Figure 3e) only in the theta band is this sufficiently common to appear in the average spectrum.

**Figure 4:**
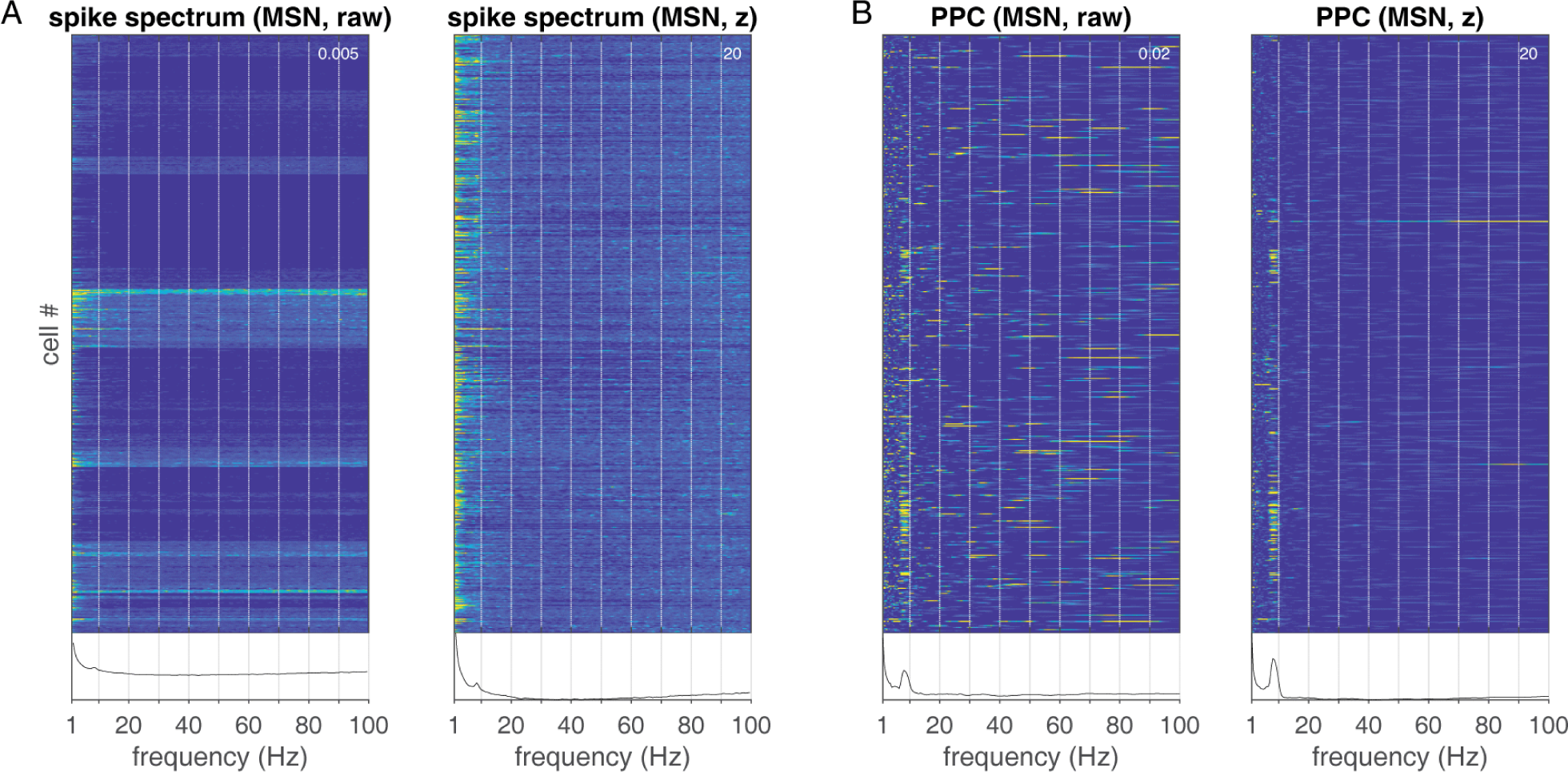
Rhythmic activity across the population of putative medium spiny neurons (MSNs, n = 579 cells) using two different measures: the spike power spectrum (**A**) which quantifies power at each frequency in the spike train, without reference to the LFP, and the pairwise phase consistency (PPC, **B**), an unbiased estimator of spike-field locking. Each row shows values for a single cell, and the inset at the bottom of each panel shows the average across all cells. Note the relatively featureless shape of the spike spectra, emphasizing low frequencies with a hint of a local increase at around 9 Hz, compared to the much richer PPC spectra, in which there is not only a clear 9 Hz peak, but in addition most other frequencies have at least some cells with increased phase locking. Left panels show raw (unnormalized) values, right panels show z-scored values against a distribution of shuffled spike times. Values in the top right corner of each panel indicate the top (yellow) end of the pseudocolor scale. Cells (rows) are ordered chronologically according to when they were acquired, such that cells recorded from each subject cluster together; this explains the apparent clustering together of similarly rhythmic cells.

The picture changes drastically when considering **spike-field locking**. For pairwise phase consistency (PPC), a measure that measures how (non-)uniformly spiking occurs across LFP phases at each frequency of interest, and for the spike-triggered average spectrum (similar, but weights spikes according to LFP power) a kaleidoscope of different phase-locking patterns is apparent across cells (Figure 4b, left). For essentially every frequency, there appear to be at least some cells that show increased phase-locking to that frequency, as indicated by the colorful streaks appearing at all possible locations on the frequency axis. Against this rich diversity of phase-locking, some overall patterns are also apparent: a clear vertical band can be seen at ∼9 Hz (theta) that also appears in the averages. Normalizing against a shuffled distribution enhances the prominence of the theta phase locking peak (Figure 4b, right).

Putative fast-spiking interneurons (FSIs) showed different patterns of rhythmic activity compared to MSNs (Figure 5, n = 154 cells analyzed that passed inclusion criteria). Like MSNs, spike spectra tended to show increased power at low frequencies (< 10 Hz), but many FSIs showed a characteristic dip in power at intermediate frequencies (10 – 30 Hz). Clear peaks in spike spectra at specific frequencies are not common, but appear less rare than in MSNs: a few examples with peaks in the low gamma range (∼50 Hz) and at theta (∼ 9Hz) can be found in the normalized spike spectra (Figure 5a, arrows in right panel). Spike-field relationships (PPC) show a clearly different pattern compared to MSNs: PPC shows the largest peak at 4–5 Hz (delta), with a substantial number of cells also locking to beta (20–30 Hz) and gamma frequencies (Figure 5b).

**Figure 5:**
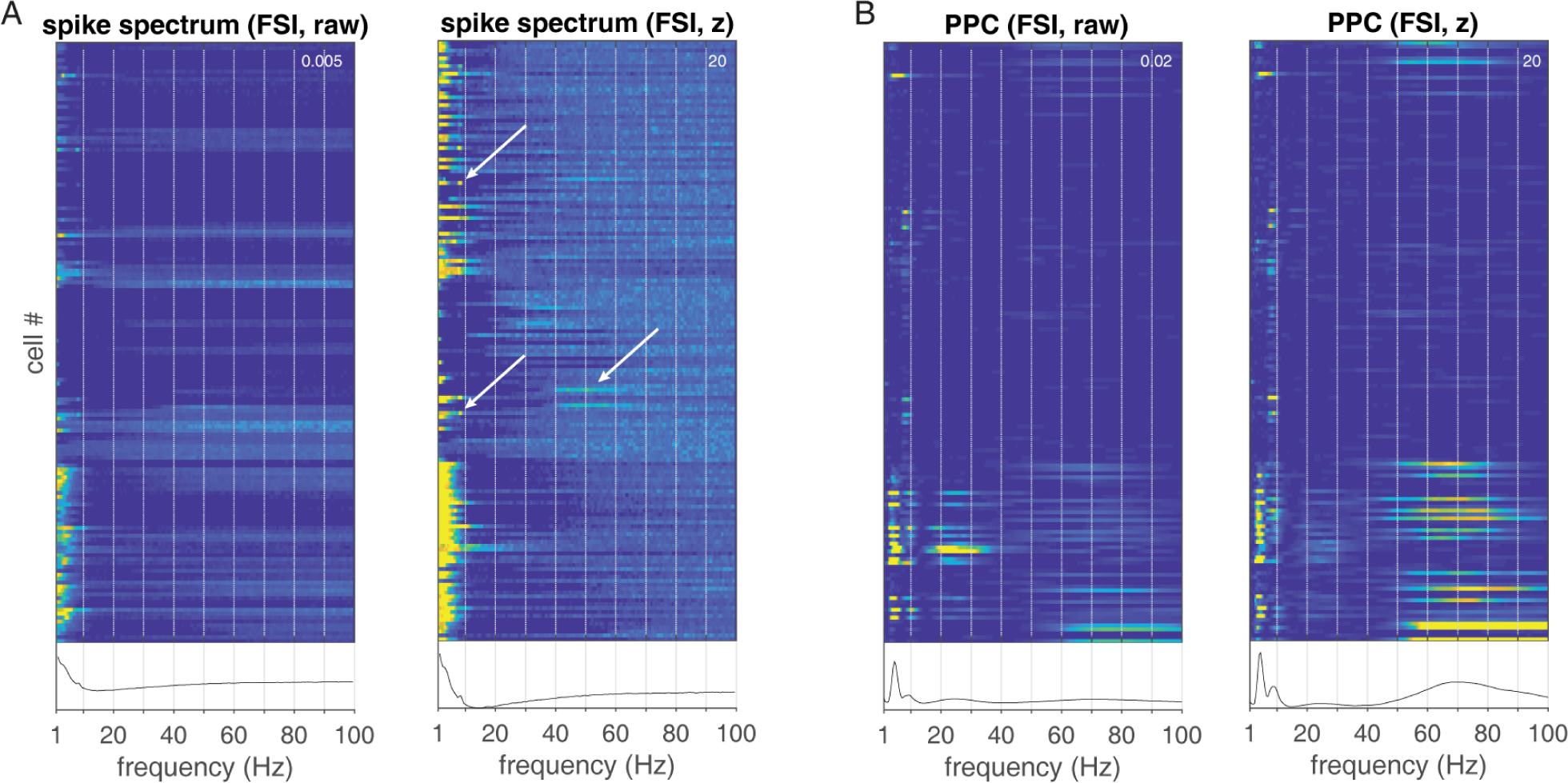
Rhythmic activity for putative fast-spiking interneurons (FSIs, n = 154), using the same layout as Figure 4. Note that the spike spectra (A) emphasize low frequencies, with barely visible peaks at delta (∼4 Hz) and theta (∼9 Hz). In contrast, spike-field relationships (B) are common in FSIs, with the delta band particularly prominent, and the normalized PPC in particular also revealing gamma-band phase locking.

To provide a more quantitative characterization of the above results, we first computed, for each frequency, what proportion of cells was significantly more rhythmic than expected by chance (at p < 0.05, uncorrected; comparison with distribution of shuffled spike trains in which spikes were randomly permuted within the 10-s analysis window). The resulting proportions (Figure 6) confirmed (1) the overall larger prevalence of spike-field locking compared to spike train rhythmicity in FSIs, (2) a theta-band peak in MSN spike spectra and phase locking, (3) elevated delta, theta and gamma-band phase locking in FSIs. Of further note is that although the increased number of spikes in typical FSIs compared to MSNs would be expected to result in a higher proportion of significant cells in FSIs, this was not found for the spike spectra, suggesting a surprising absence of rhythmic spiking in FSIs despite widespread phase locking (consistent with desynchronization in FSIs in experimental and modeling work; Berke 2008; Hjorth et al. 2009).

**Figure 6:**
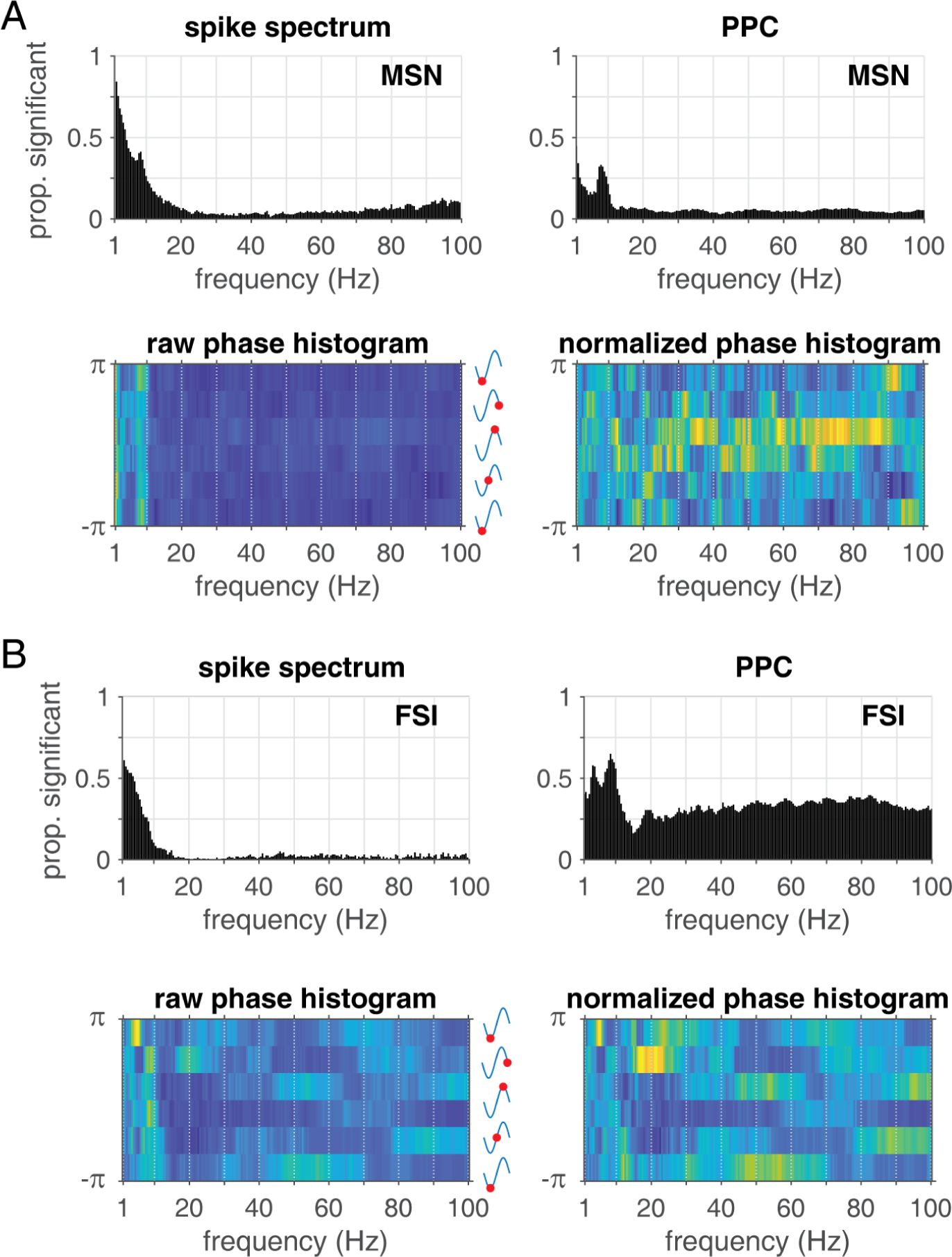
**A:** Proportions of putative MSNs with significant spike spectrum power (top left) and significant spike-field locking (top right). Bottom row shows histograms of preferred firing phases for each LFP frequency of interest. Color indicates the number of cells preferring to fire at each of six phase bins; if each neuron individually preferred a random phase (no population phase preference) then these counts would be uniform; by contrast, a population phase preference would manifest as a difference in counts across phase bins. Both the raw (left, the sum of each column corresponds to the number of cells with significant phase locking at that frequency) and normalized (right, counts in each column divided by the number of cells) histograms show some evidence of non-uniform phase preferences, indicating some temporal coordination (synchrony) across the population. **B**: Same as A, but for putative FSIs. Note the overall higher prevalence of significant spike-field relationships, but not spike spectra, compared to MSNs.

A related property of interest is the preferred firing LFP *phase* of ventral striatal neurons. The distribution of preferred phases across the population of neurons can provide important clues about a number of issues, such as (1) whether there a preferred phase across the population, which would indicate temporal coordination (synchrony), (2) whether there are systematic preferred phase differences between cell types, indicative of local interactions (e.g. inhibitory interneurons firing before projection neurons), and (3) relationships between anatomically related structures (e.g. preferred phases may be consistent or inconsistent with the preferred phase of another brain area). To characterize preferred phases across the population, we first computed, for each frequency, a histogram of preferred phases (Figure 6, bottom row; note that for each frequency, only neurons with significant phase locking at that frequency are included). In the raw histogram (left panel), each column sums to the total number of significantly phase-locked neurons for that frequency (as given by the histogram in Figure 6, top row). An absence of phase preference across all significantly phase-locking neurons would appear as constant values within a column (frequency); conversely, a clear-phase preference would appear as a peak within a column. As Figure 6 indicates, most frequencies showed a moderate amount of non-uniform phase distributions, suggesting some amount of coordination across the population.

The above histograms ignored the strength of phase locking, by pooling all neurons with significant phase locking and treating them equally. However, the preferred phase of neurons with strong phase locking is likely more meaningful than that of neurons with a barely significant phase preference. The polar plots in Figure 7 highlight that for some frequency bands, the most strongly phase-locked neurons maintain a consistent phase (e.g. delta-band for FSIs and MSNs, left column) whereas for other frequency bands, there is a much weaker population preference (e.g. theta-band, perhaps related to the known tendency of such neurons to phase-precess, van der Meer and Redish 2011).

**Figure 7:**
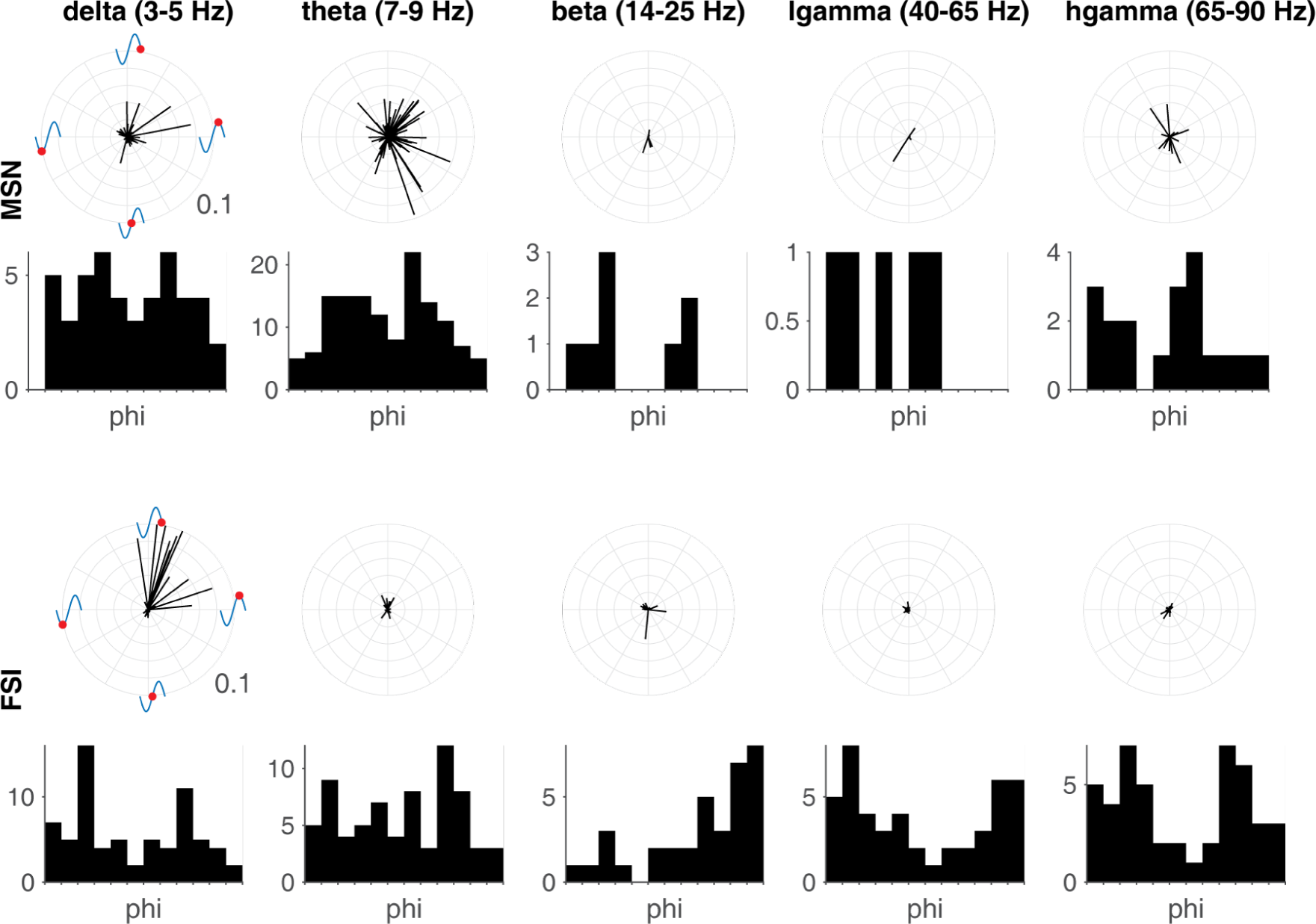
Polar plots showing both phase preference (angle) and strength of phase locking (length) for each cell with significant phase locking in each of five frequency bands of interest. Each line corresponds to a single cell. Phase histograms within each frequency band are also shown; these are constructed in the same way as the raw phase histograms in Figure 6, except that all frequencies within each band are averaged, and more phase bins are used (12 instead of 6). Top panel shows MSN data, bottom panel FSI data. Note that some frequency bands show clearly non-uniform phase distributions (e.g. all frequency bands except perhaps theta for FSIs) whereas others don’t show a clear phase preference (e.g. delta band for MSNs) or don’t have enough significantly phase-locked cells to make a determination (beta band and up for MSNs). All polar plots use the same scale, with the outermost ring indicating a PPC value of 0.1.

### Relationships between different aspects of rhythmic activity

In principle, rhythmic spiking and phase locking can be completely independent: a given cell may spike metronomically at some frequency but without any relationship to the LFP, leading to the absence of a phase preference. Similarly, a cell may be perfectly phase locked to say, LFP theta, in the sense that when it does spike it does so at a specific phase. But if this cell only fires every few seconds, and/or LFP theta deviates from a perfectly constant frequency, this phase preference would not translate into a spike spectrum peak. Thus, we can ask how correlated, across cells, spike spectrum power and phase locking measures are. The simplest version of this analysis simply correlates, for each frequency, the values across all cells for one measure (e.g. spike spectrum power) with another (e.g. PPC). The results of this analysis are shown in the lower quadrant of Figure 8. As expected, PPC and STA power are highly correlated overall, with noticeable peaks in the theta, beta, and low-gamma ranges (bottom row, center panel). In comparison, spike spectrum power is noticeably less correlated with PPC and STA (left column, bottom two panels), with only theta and beta frequencies showing a moderate relationship. This confirms the impression from Figures 3b-d and 4–5 that for many frequencies there is not an obvious relationship between spike spectrum power and spike-field locking.

**Figure 8:**
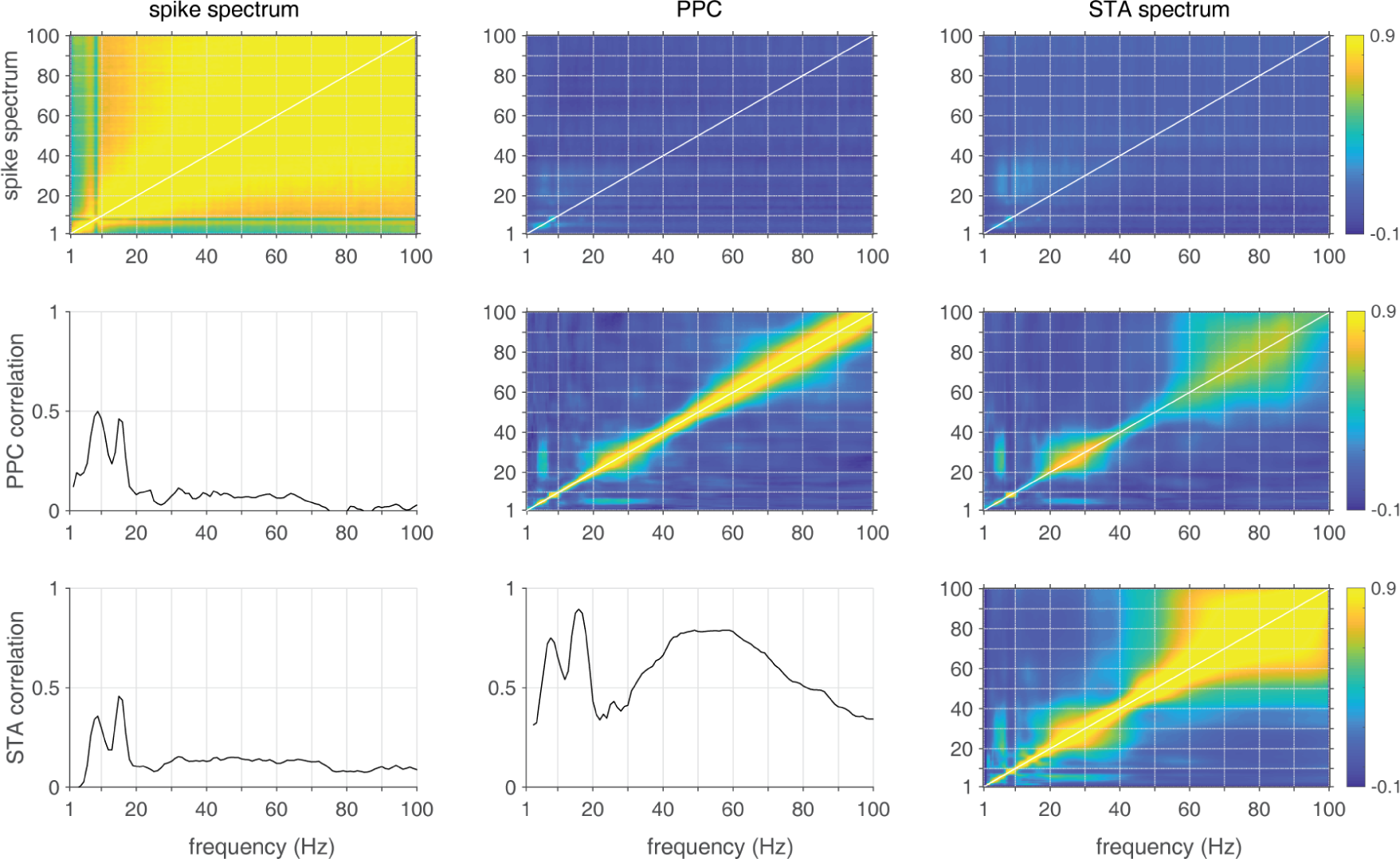
Correlations between spike spectrum power (SS), pairwise phase consistency (PPC) and spike-triggered average (STA) spectrum measures. The *lower quadrant* of panels shows correlations between these measures for the same frequency: for instance, the middle panel in the left column shows the correlation (across all cells, one frequency at a time) between spike spectrum power and PPC. Note the comparatively higher correlation between PPC and STA power (center column, bottom panel) compared to the correlations between spike spectrum power and the spike-field measures (left column, bottom two panels). The *upper quadrant* of panels shows the full correlation matrices across frequencies. The diagonal (white line) of these matrices is identical to the correlations in the lower quadrant. These matrices illustrate relationships of the type visible, for instance, in the center panel: cells that tend to phase-lock to 20-40 Hz also tend to phase lock to ∼ 4 Hz, as illustrated by the off-diagonal patches of increased correlation for these frequencies.

### Predicting ventral striatal spike times based on LFP features

The analyses so far have shown that a substantial percentage of vStr neurons shows phase-locking to one or more frequencies in the LFP. Although the above analyses are useful in demonstrating widespread spike-field relationships across the population of ventral striatal neurons, the PPC and STA spectrum measures have limitations. First, it is hard to know “how important” of a contribution this phase locking is to all other factors that are related to spike timing, such as tuning for task variables, the cell’s autocorrelation, and so on. An extreme version of this issue is that if a cell has a tendency to burst at a given frequency, some phase locking may result, but LFP features would provide no new information above and beyond what could already be predicted based on the burst properties. Second, want to find out how much oscillations in other brain areas contribute to the prediction of vStr spike timing.

To address these limitations, we fit generalized linear models (GLMs) to ventral striatal spike trains to quantify the added predictive value of the ventral striatal LFP (above and beyond spiking and task variables) and the added predictive value of a LFP from an anatomically connected structure, the hippocampus. To quantify the fit of these different models, we used cross-validation: across different splits of the available data for each cell, models were fit to one half of the data, and then applied to the withheld data to yield error measures for each cell and model. The first overall question addressed with this approach is simply, for how many vStr cells does knowledge of LFP phase improve spike timing prediction, above and beyond predictions made from a number of influential predictors such as the cell’s autocorrelation and tuning for task variables (see *Materials and Methods* for details)? As shown in Figure 9a, the vast majority of cells (89.6%) benefited from the addition of LFP features. Comparing the contributions of different predictors reveals that the cell conditional intensity function (“ci”) is the most influential predictor overall, with time-to-reward (“tt”) and position on the track (“li”) the next most important (Figure 9b-d). Then, however, LFP phase at various frequencies consistently contribute, led by delta and theta, outperforming running speed (“sp”). Thus, this GLM analysis further confirms the prevalence of spike-field relationships shown in the previous analyses with a more stringent criterion, and reveals the predictive power of the LFP to be comparable to that of previously established task- and behavior-related predictors.

**Figure 9:**
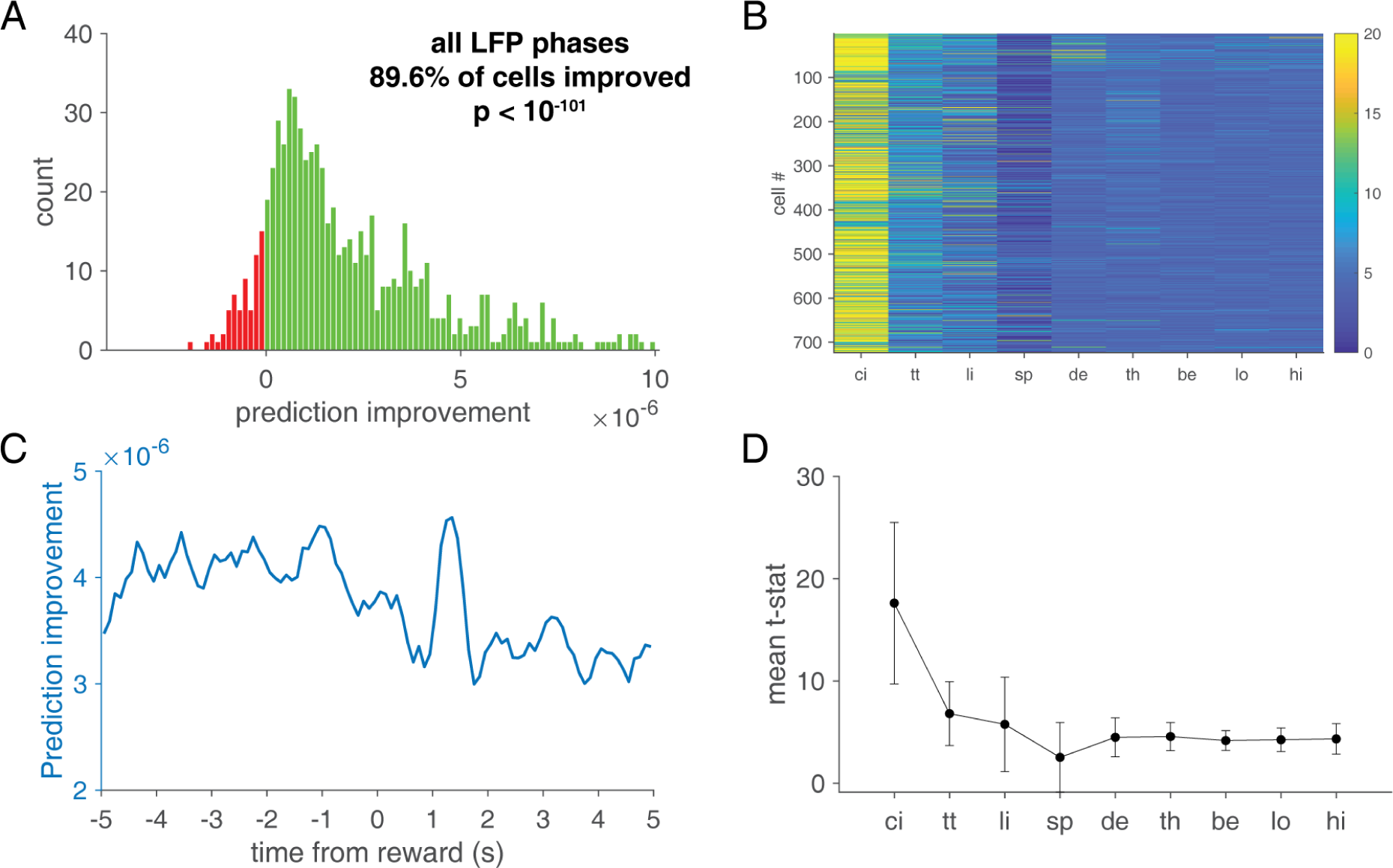
LFP phase improves cross-validated prediction of ventral striatal spike timing. **A**: Overall model comparison between a baseline model (without any LFP features) and a model with all LFP features included (the phase of each of the five frequency bands). Positive numbers (indicated in green) indicate better model fit for the LFP model compared to baseline, red bars indicate worse model fit. **B**: Relative contribution for each cell (rows) of different predictors (ci: conditional intensity function, tt: time to reward, li: linearized position, sp: running speed, and phase for each frequency band of interest (delta, theta, beta, low-gamma and high-gamma); the average t-statistic for each predictor is shown in panel D. **C**: Model prediction improvement plotted as a function of time from reward delivery. Overall, LFP features provided more improvement before reward delivery (negative time from reward) compared to after reward delivery, other than a peak around 1.5s (the time of first contact with reward pellets).

Unlike typical cross-validation approaches that simply sum all errors across the testing set to yield one final error measure, we also tracked errors according to when in the task, relative to reward delivery, they occurred (see Figure 2 for a schematic illustration of this approach). These errors can then be visualized as a peri-event average around the time of reward delivery (Figure 9c), showing that the improvement in model prediction is not constant over time. Furthermore, we can break out the contribution of each LFP frequency band to reveal its contribution as a function of time (Figure 10). For the delta and theta bands in particular, LFP phase was more predictive before, rather than after, reward delivery. In comparison, low gamma phase was more informative following reward, rather than before; as can be seen from the close correspondence with LFP low-gamma amplitude (red line in top right panel in Figure 10) this pattern results from a simple increase in signal-to-noise in the low-gamma band LFP. However, in general it was not the case that the contribution of LFP phase to model performance is not determined simply by the amplitude of the corresponding oscillation; this can be seen by comparing the model improvement (blue lines) with the amplitude (red lines) for the delta and theta frequency bands. This dissociation shows that dynamic changes in spike-field relationships in the ventral striatum are not an artifact of variations in the ability to accurately measure LFP phase.

**Figure 10:**
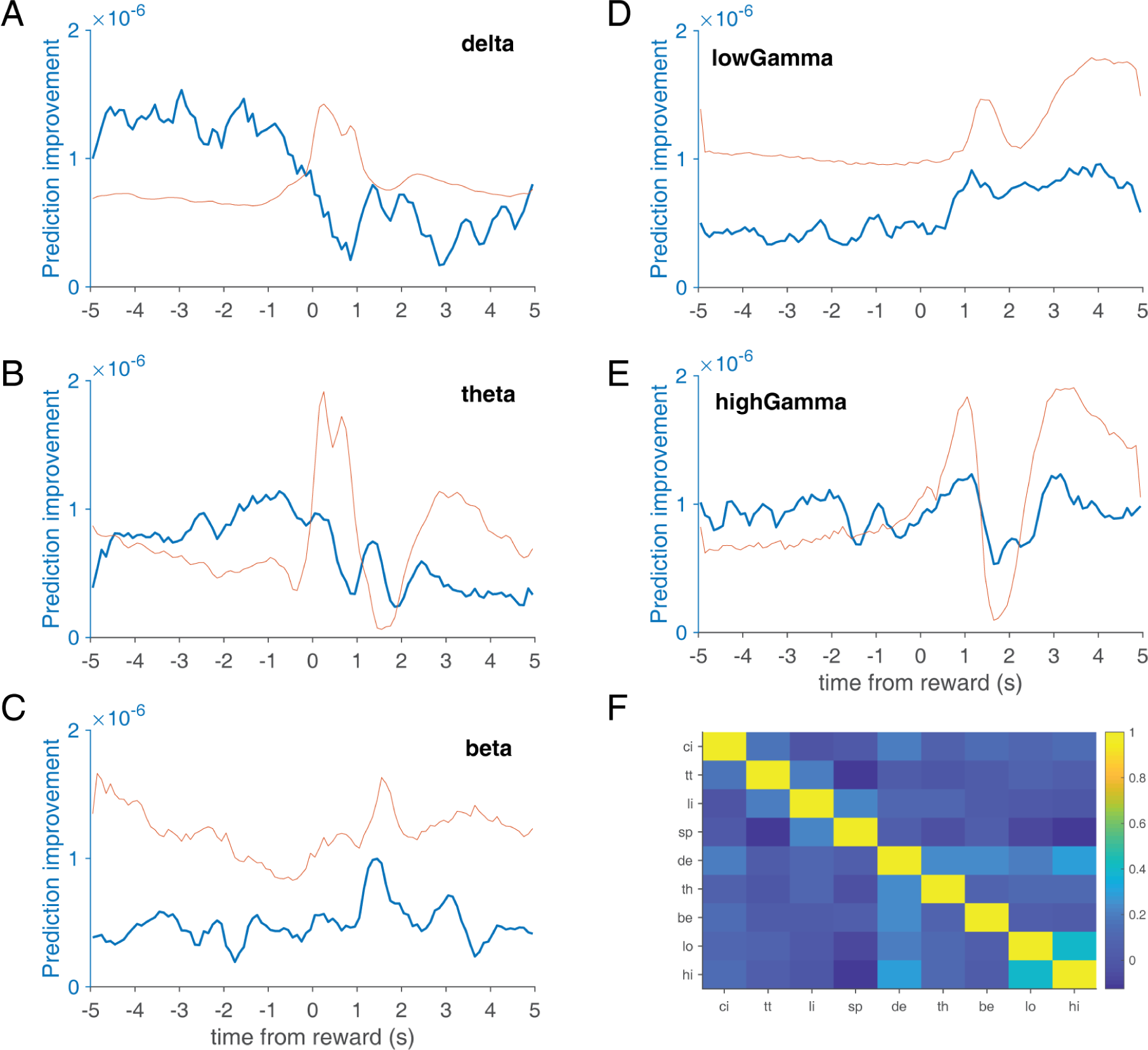
**A-E:** Model improvement relative to baseline (blue) and LFP envelope (red) for each LFP frequency band of interest as a function of time relative to reward. Note that for delta and theta bands in particular, LFP phase contributes more to model fit before, rather than after, reward delivery in a manner that cannot be explained by differences in the LFP envelope (red line). In contrast, in the low-gamma band (top right panel) LFP phase contributes in a manner consistent with the changes in amplitude. **F**: Correlation matrix between all predictors in the full model.

A final application of our GLM approach we consider is to reveal the contribution of a LFP from *different* brain area: in this case, the hippocampus (HC), which is one of several limbic brain structures that project monosynaptically to the vStr (van der Meer et al., 2014). Changes in the ability of such a distal LFP to predict local spike times are a compelling way to operationalize the idea of effective connectivity, that is, the extent to which oscillatory activity in one brain area contributes to activity in another (Wong et al., 2016). In particular, this approach takes an important step beyond showing changes in LFP relationships such as coherence, which has been previously shown to increase between HC and vStr as animals approach reward sites (van der Meer and Redish, 2011; Lansink et al., 2016; Sjulson et al., 2018) but which cannot reveal changes in spiking activity. Our GLM approach enables us to ask first, whether the HC LFP contributes to prediction of vStr spiking overall, and second, where during the task that contribution is strongest. Figure 11a shows the overall contribution, which shows a clear improvement for the majority of cells (note the number of cells here is different to the earlier analyses because this is taken from subjects with recordings from both vStr and HC, whereas the earlier analysis was vStr only; the lower number of cells means we combined putative cell types). The distribution of this improvement across the task (Figure 11b) demonstrates that on average, the contribution of HC theta phase to vStr spike timing peaks around the time of reward delivery. As with the previous analyses, changes in model improvement did not simply track theta oscillation amplitude (red line). In any case, this analysis illustrate the versatility of the GLM approach in enabling a spike-based measure of functional and effective connectivity applied to the ventral striatum.

**Figure 11:**
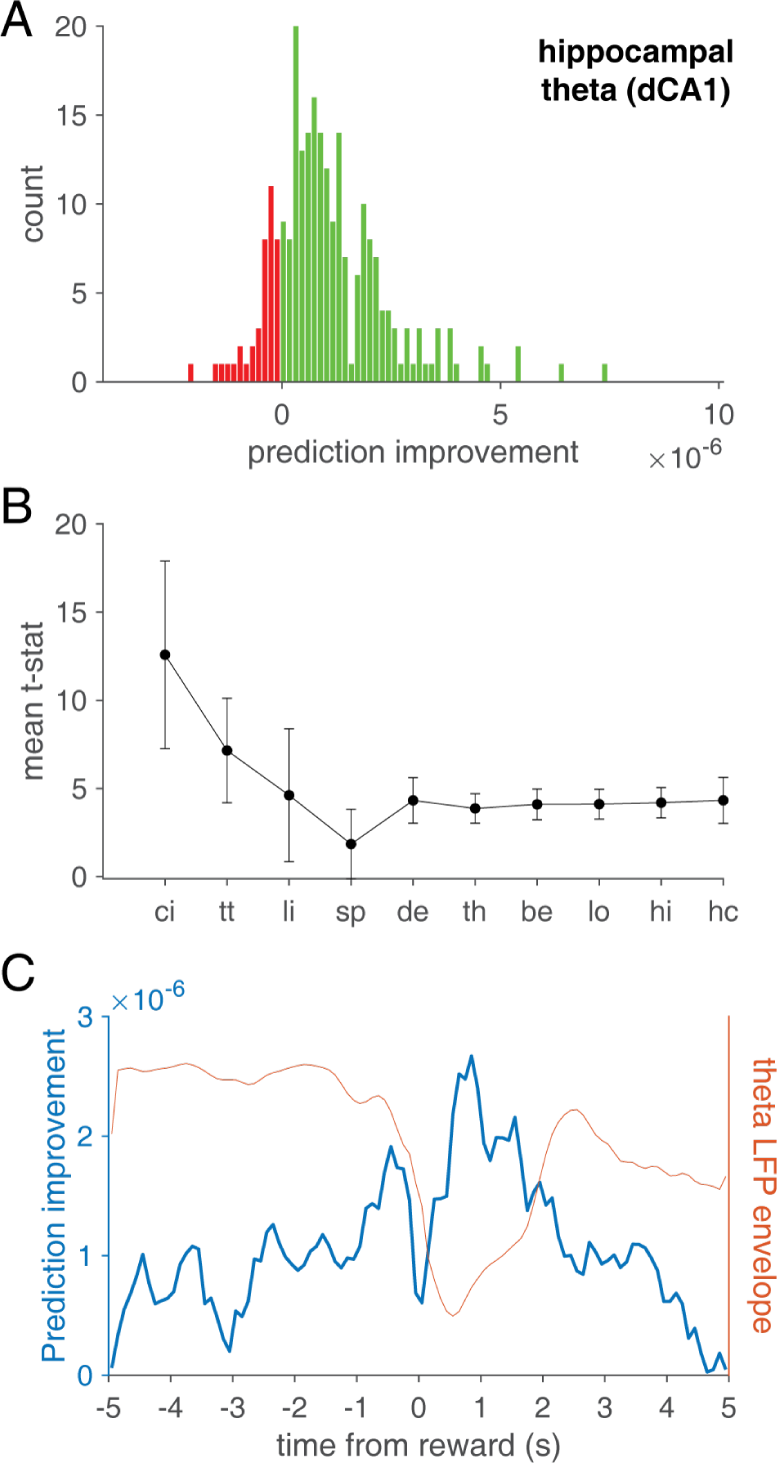
Hippocampal LFP phase improves prediction of ventral striatal spike timing. **A**: Overall model comparison between the baseline model (containing the full set of predictors used earlier, including the phases at each frequency band obtained from the ventral striatal LFP) and the baseline model with hippocampal theta phase added. The hippocampal theta model significantly improved model fit overall. **B**: Mean t-statistics for all predictors in the model. As before, the cell autocorrelation is the best predictor overall, and all LFP features contribute approximately equally, although hippocampal theta (hc) was numerically better than any ventral striatal LFP feature. **C**: Model prediction improvement (blue line) as a function of time from reward was largest around the time of reward delivery. This pattern was not explained simply by theta amplitude being largest at that time, as indicated by the envelope (red line) which showed an opposite pattern.

## Discussion

### Summary

We have shown that spike-field relationships are pervasive in ventral striatal neurons, with different MSNs and FSIs showing phase locking in essentially every frequency band. MSNs tended to favor theta-band phase preferences, while FSIs favored delta and gamma overall. Spike train rhythmicity without reference to the LFP showed more modest frequency content, and was generally independent of phase-locking except for theta and beta bands – that is, a theta phase-locked neuron is likely to spike at theta frequency, but a gamma phase-locked neuron is no more or less likely to spike at gamma frequency. Setting a high bar for phase-locking, we used generalized linear models (GLMs) to show that even after the influence of cell and task variables is accounted for, LFP phase improves spike timing prediction for the vast majority of neurons. Interestingly, the contribution of the LFP to this prediction is dynamic, which we use to (1) rule out simple changes in LFP oscillation amplitude as an explanation, and (2) enable new measures of functional coupling by using the LFP from an input structure (hippocampus) to predict local spiking.

These findings inform the larger project of establishing how rhythmic activity in the ventral striatum can be leveraged for a better understanding of this widely studied brain area’s structure and function. We believe this is a productive approach for a number of reasons. First, an oscillatory perspective on brain activity has been tremendously successful in a number of anatomically related regions, such as navigation and memory in the hippocampus (Buzsáki, 2002; Burgess and O’Keefe, 2011; Colgin, 2016) and perception in the cortex (Cardin et al., 2009; Bastos et al., 2015; VanRullen, 2016). Second, many theories of ventral striatum function emphasize the importance of switching between multiple convergent inputs (Finch, 1996; Grace, 2000; Gruber et al., 2009), which is exactly the type of operation that rhythmic fluctuations in neural activity have been proposed to contribute to (Akam and Kullmann, 2010; Fries, 2015). Finally, oscillations are well positioned to connect neural phenomena at different spatial and temporal scales, including single-cell properties such as resonance, microcircuit interactions, and long-range communication between different brain regions (Berke, 2005; Buzsáki, 2006). These systems-level interactions are increasingly thought to underlie the most mysterious aspects of cognition and its dysfunction.

### Relationship to previous work

Several previous studies in behaving animals have examined spike-field relationships in the ventral striatum. However, these studies have generally focused only on a specific frequency band (e.g. gamma, van der Meer and Redish 2009, Kalenscher et al. 2010; beta and gamma, Howe et al. 2011; theta, van der Meer and Redish 2011) or cell type (e.g. FSIs only, (van der Meer and Redish, 2009); MSNs only, (Kalenscher et al., 2010). Because these studies used different behavioral conditions during which data were acquired, as well as different analysis methods to identify and quantify phase locking, a comprehensive picture across frequency bands and cell types has been lacking. Moreover, to our knowledge no previous work has examined spike train rhythmicity without reference to the LFP. This is an important issue because in general, the LFP contains contributions from a number of sources which may only be indirectly linked to local spiking (Buzsáki et al., 2012; Wilson et al., 2018; Pesaran et al., 2018). For the vStr in particular, the connection between the LFP and local spiking may be even more indirect than is typical, because the non-layered geometry of the striatum implies the ventral striatal LFP is dominated by volume-conducted components (Carmichael et al., 2017). Thus, oscillations in the vStr LFP and local spiking may range from strongly related to essentially disconnected.

Comparing the results from our comprehensive approach with previous work highlights areas of agreement as well as a number of novel observations. Among the more striking observations is the finding that phase locking is widespread across the population of vStr neurons (about 90% of neurons showing a relationship to at least one frequency band, as measured by (1) improved spike time prediction based on LFP phase in generalized linear models, and (2) the diverse regions in frequency space highlighted by the PPC plots in Figures 4-5). This prevalence of spike-field relationships across frequencies had not been apparent from isolated reports of e.g. ∼10% gamma-phase locking neurons (Kalenscher et al., 2010) or ∼15% theta-phase preferring neurons (van der Meer and Redish, 2011). We consistently found delta phase locking, in line with data from anesthetized animals (Leung and Yim, 1993) and isolated examples visible in figures making a different point (Berke, 2005; van der Meer and Redish, 2009), which fits with reports of a widespread 4 Hz oscillation in the limbic system (Fujisawa and Buzsáki, 2011; Karalis and Sirota, 2018).

A related new contribution in this study is the systematic investigation of preferred phases, which tends to be an overlooked issue in studies of rhythmic activity. At the population level, there is a major difference between (a) neurons having a uniform distribution of preferred phases, and (b) coordination between preferred phases across neurons (synchrony); merely reporting that a given percentage of neurons shows significant phase locking does not distinguish between these possibilities. Importantly, in the latter case, the LFP is much more informative about the state of the population than in the former. We found that preferred phases of MSNs in particular, but to some extent also FSIs, could be surprisingly non-uniform. This may be in part due to the fact that our 10-s window contained multiple distinct network activity states (reward approach/expectation, movement and non-movement, reward receipt and consumption) but could also be related to the emerging idea that local connectivity between striatal neurons (such as gap junctions between FSIs) paradoxically may facilitate de-correlation rather than synchronization (Hjorth et al., 2009; Gage et al., 2010). Similarly, excessive population synchronization may in fact indicate pathological activity such as occurs in advanced Parkinson’s Disease (Jenkinson and Brown, 2011). Further work comparing preferred phase in a time-resolved manner, perhaps during different task components as well as rest/sleep states, can shed some more light on this matter.

A final innovative aspect of this work is the application of generalized linear models (GLMs) to quantify spike-field relationships in the vStr. At the most basic level, this approach sets a high bar for spike-field locking by incorporating possible covariates; it also allows for systematic comparison of the relative contribution of phase locking to predicting spike times compared to other (task) variables. In this respect, the overall contribution of LFP phases to spike timing was on par with known task variables such as time to reward delivery and position on a maze. By retaining the time-resolved model improvement for different GLMs, we were able to quantify the contribution of LFP variables not only overall, but as a function of time. This time-resolved analysis shows that the contribution of LFP variables varies in a manner that is not simply predicted by changes in the amplitude of the LFP. This dynamic spike-field locking may be expected if the LFP itself contains large volume-conducted components (Carmichael et al., 2017), because the source(s) of the LFP may be more or less effective in contributing to vStr spiking depending on context.

### Limitations and future work

As highlighted above, we only analyzed a 10s window around a salient task event (reward delivery); we did not break down this window into more specific components, nor did we analyze rest/sleep, where rhythmic activity including spike-field locking may well be different. Comparisons between different task components and behavioral states could help disentangle the relative contributions of cell-specific mechanisms (such as particular ion channel distributions) on the one hand, and network-level emergent properties on the other. Contributions of single cell channels may be expected to be relatively stable such that there may be “oscillatory fingerprints” of specific cells, which may get different inputs, project to different places, and may relate to what task variables are coded.

This data set used extracellular recordings from chronically implanted animals, which comes with a number of standard caveats: our cell classification was not verified with e.g. single cell morphology or I/V curves, but based on waveform and firing characteristics only. In addition, although we estimated approximate recording locations for this same data set in earlier work (van der Meer and Redish, 2009, 2011) we do not have more detailed information on whether a given cell was recorded in say, the core or shell of the nucleus accumbens. Inspection of the figures in this study suggests that there are differences in rhythmic activity between subjects (e.g. Figure 4-5) which may be attributable to differences in recording location. Future work using systematically arranged recording arrays and/or tagging of specific cell types or projection targets with optogenetic tools will be able to test this idea more thoroughly.

Finally, returning to the overall questions raised in the introduction, what does this detailed view of vStr rhythmic spiking provided by this study tell us about how important oscillations are for the function of this structure? Now that we have a comprehensive baseline of such activity, comparisons with animals or strains that model aspects of human disease are an obvious next step, as are interactions with neuromodulators such as dopamine. In addition, the widespread nature of oscillatory activity in the vStr keeps alive the notion that coherence with anatomically related areas (or the absence of it) may have functional consequences. This idea could be tested formally by stimulating an input to the vStr at different phases of the ongoing LFP, and determining whether these phases are associated with a different probability or magnitude of a response (Cardin et al., 2009; Womelsdorf et al., 2012). The ubiquity of oscillatory inputs in the vStr and the convergence of multiple such inputs onto single neurons and local circuits suggests that such studies will be informative.

## Acknowledgments

These data were recorded in the lab of A. David Redish at the University of Minnesota.

## Author contributions

MvdM performed experiments and pre-processed the data. JMG, JEC and MvdM wrote analysis code and performed data analysis. MvdM wrote the paper with comments from JMG and JEC.

## Footnotes

1 Note that PPC is an estimate of the *square* of the phase-locked value (mean vector length across spike phases), such that a PPC of 0.01 corresponds to a mean vector length of approximately 0.1.

2 Note that it is possible in principle to have rhythmic spiking without phase locking; and vice versa.

